# A recombinant rotavirus harboring a spike protein with a heterologous peptide reveals a novel role of VP4 in viroplasm stability

**DOI:** 10.1101/2021.05.13.444118

**Authors:** Guido Papa, Janine Vetter, Michael Seyffert, Kapila Gunasekera, Giuditta De Lorenzo, Mahesa Wiesendanger, Elisabeth M. Schraner, Jean-Louis Reymond, Cornel Fraefel, Oscar R. Burrone, Catherine Eichwald

## Abstract

The rotavirus (RV) VP4 spike protrudes as a trimeric structure from the five-fold axes of the virion triple-layer. Infectious RV particles need to be proteolytically cleaved in VP4 into two subunits, VP8* and VP5*, constituting both the distal part and central body of the virus spike. Modification of VP4 has been challenging as it is involved in biological process including the interaction with sialic acid and integrins, cell tropism and hemagglutinin activity. Here, we engineered a loop at position K145-G150 in the lectin domain of the VP8* subunit to harbor a small <u>b</u>iotin <u>a</u>cceptor <u>p</u>eptide (BAP) tag and rescued viable viral particles using RV reverse genetics system. This rRV/VP4-BAP internalizes, replicates, and generates virus progeny, demonstrating that the VP4 spike of RV particles can be genetically manipulated by the incorporation of at least 15 exogenous amino acids. Although, VP4-BAP had a similar distribution as VP4 in infected cells by localizing in the cytoskeleton and surrounding viroplasms. However, compared to wild-type RV, rRV/VP4-BAP featured a reduced replication fitness and impaired viroplasm stability. Upon treatment of viroplasms with 1,6-hexanediol, a drug disrupting liquid-liquid phase-separated condensates, the kinetic of rRV/VP4-BAP viroplasm recovery was delayed, and their size and numbers reduced when compared to viroplasms of wild type RV. Moreover, siRNA silencing of VP4 expression in RV strain SA11 showed similar recovery patterns as rRV/VP4-BAP, revealing a novel function of VP4 in viroplasm stability.

**IMPORTANCE:** The rotavirus (RV) spike protein, VP4, has a relevant role in several steps involving virion internalization. The strategic position of VP4 in the virion resulted in a challenge for the addition of an exogenous peptide producing infectious particles. The identification of a specific loop in position K145-G150 in the VP8* subunit of VP4 allowed the rescue by RV reverse genetics of a recombinant RV harboring VP4 containing a 15 amino acids tag. This study demonstrates this recombinant virus has similar replication properties as a wild-type virus. Moreover, we also discovered that VP4 is necessary for the assembly and stabilization of the cytosolic replication compartments, the viroplasms, demonstrating a novel role of this protein in the RV life cycle.

## INTRODUCTION

Rotavirus (RV) is the primary etiological agent responsible for severe gastroenteritis and dehydration in infants and young children worldwide (1) as well as young animals such as piglets, calves, and poultries, thus representing a negative economic impact on livestock (2–4). RV virions are non-enveloped particles composed of three concentric layers. The virus core-shell encloses eleven double-stranded RNA (dsRNA) genome segments and twelve copies of the structural proteins VP1 (the RNA-dependent RNA polymerase) and the guanyl-methyltransferase VP3 (5, 6). The icosahedral core-shell (T=1, symmetry) is composed of twelve decamers of VP2 and surrounded by 260 trimers of the abundant structural protein VP6, constituting the transcriptionally active double-layered particles (DLPs) (7, 8). On top of each VP6 trimer stands a trimer of the glycoprotein VP7, the main building component of the outer layer of the virion that also shows an icosahedral symmetry (T=13). At each of the 5-fold axes of the outer layer, the VP4 spike protein is anchored in a trimeric conformation, although adopting a dimeric appearance when visualized from the above capsid surface (9–12). Infectious RV virions, also named triple-layered particles (TLPs), rely on the cleavage of the VP4 spike protein by a trypsin-like enzyme found in the intestinal tract (13). The proteolytic cleavage of VP4 (88 kDa) entails two main products, VP8* (28 kDa, amino acids 1-247) and VP5* (60 kDa, amino acids 248-776) that remain non-covalently associated with the virion (14, 15). VP8* and a significant portion of VP5*, VP5CT (amino acids 246-477), as analyzed by crystallography, constitute the distal globular density and the central body of the spike, respectively (16, 17). The VP8* subunit has several functions such as haemagglutinin activity (18), involvement in the binding to sialic acid (17), and a determinant role in virus tropism, while VP5* has been implicated in the interaction with integrins (19–21). Interestingly, VP4 is not only involved in RV tropism, attachment, neutralization, and entry into host cells, but has also been shown to play an essential role in virion morphogenesis. During virus internalization, VP4 was shown to bind to the small GTPase Rab5 and PRA1 within early endosomes (22), to directly activate the heat-shock protein 70 (23, 24) and to bind to the actin-binding protein drebrin (25) followed by association to the microtubules and actin cytoskeletons (26–31).

Specific *in vivo* biotinylation of cellular targets can be achieved by adding a small <u>b</u>iotin <u>a</u>cceptor <u>p</u>eptide tag (BAP) (15 amino acids) to the protein of interest and co-expressing with the *Escherichia coli*-derived biotin ligase, BirA (32). This enzyme covalently links a single biotin molecule to the unique lysine within the BAP tag (33, 34). This methodology is a powerful biotechnological tool for versatile applications, such as identifying highly complex interactomes (35, 36), and permits batch protein and subviral particle (33, 34) refinement at high purity and in physiological conditions. An example is the incorporation of a BAP tag in RV VP6, allowing preparation and purification of replication-competent DLPs (33).

Here, we describe the generation of a recombinant RV (rRV) harboring a genetically modified genome segment 4 (gs4) encoding the structural protein VP4 with an in-frame inserted BAP tag in an external loop of the VP8* subunit. The biotinylated rRV/VP4-BAP can infect, replicate and generate virus progeny. Moreover, it revealed a novel function of VP4 associated to viroplasm assembly and stability.

## RESULTS

### Production and characterization of recombinant rotavirus expressing VP4-BAP protein

As VP4 is the main structural RV protein involved in host cell tropism, attachment, and internalization, we addressed the question whether it could be engineered by incorporating a peptidic tag within its coding sequence without compromising its structural and functional properties. To test this hypothesis, we used the previously published crystal structure of simian RRV VP4 (10) to identify four different loops localized in the lectin domain (amino acids 65-224) of the VP8* subunit and then inserted a BAP tag (33, 34) in the corresponding loops of the simian strain SA11. As depicted in **Fig. 1A**, the selected amino acid regions for the BAP tag insertions were T96-R101, E109-S114, N132-Q137, and K145-G150 of VP4 of strain SA11, assigned with colors blue, orange, pink, and green, respectively. The biotinylation of these BAP-tagged VP4 proteins was then analyzed in total cell lysates in a Western blot-retardation assay (WB-ra) (34). Each of these constructs, driven by a T7 promoter, was co-transfected with a DNA plasmid encoding the cytosolically localized enzyme BirA (cyt-BirA) into MA104 cells, which were also infected with a recombinant T7 RNA polymerase vaccinia virus to allow cytosolic transcription (37). It has been noticed that VP4 was not expressed when transcribed using a nuclear promoter, probably due to mRNA splicing of the VP4 transcript. As shown in **Fig. 1B**, the four VP4-BAP variants (**Fig. 1B, lanes 3-10**), but not the wild type (wt) VP4 (**Fig. 1B, lanes 1 and 2**), were fully biotinylated. Of note, the band detected above the VP4-BAP band corresponds to a phosphorylation form of the protein only present in transfected cells but not in RV-infected cells, as demonstrated by the λ-phosphatase treatment (**Fig. S1A and S1B**). Taken together, the results indicate that the expression and stability of the different VP4 protein mutants were not affected by the location of the inserted BAP tag.

**Figure 1.**
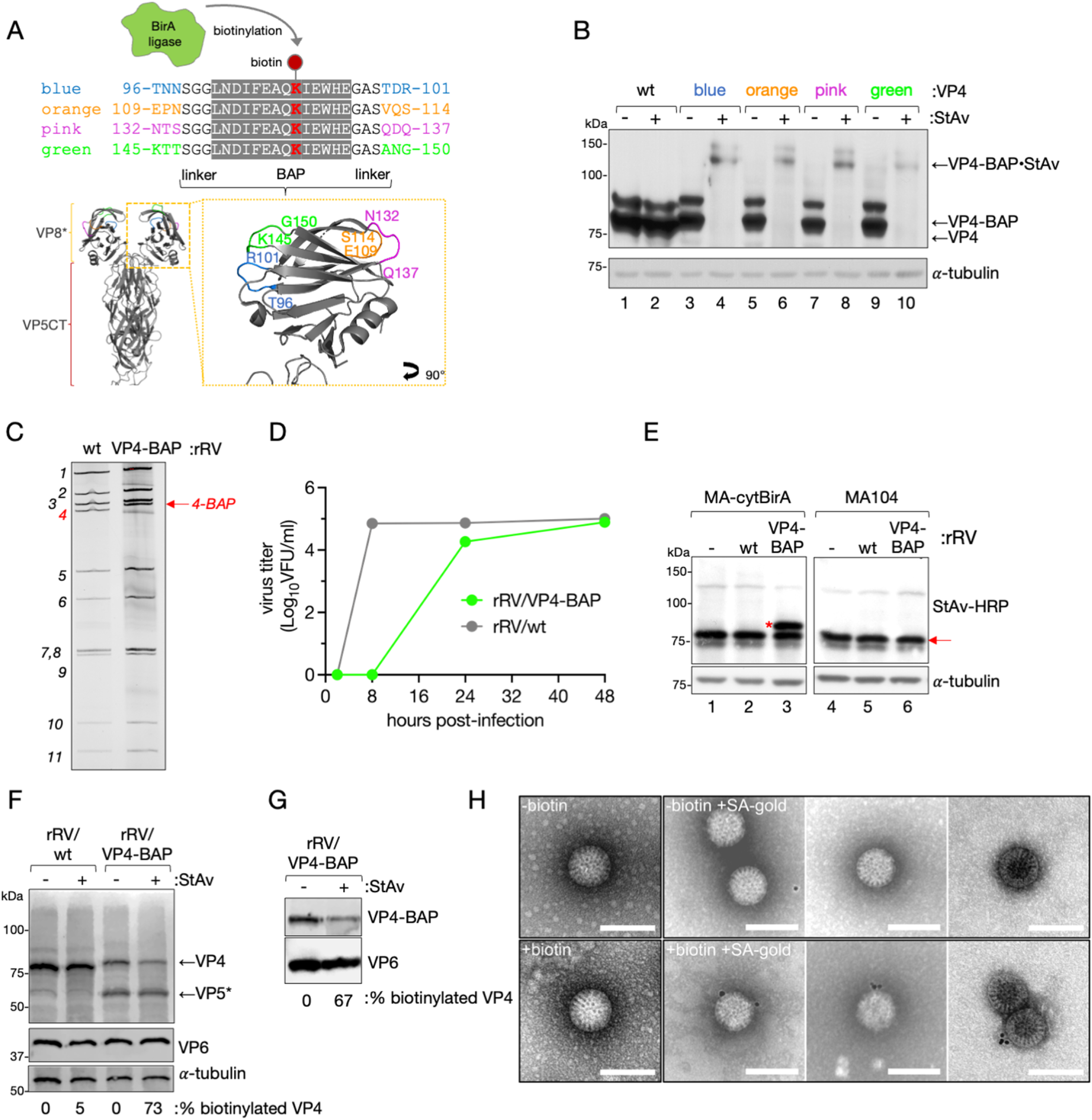
Generation of VP4-BAP tagged recombinant rotavirus. **(A)** Schematic representation of BAP tag inserted in lectin domain loops of the VP8* subunit of VP4 from RV simian strain SA11(GenBank: X14204.1). The lysine (K, red) indicates the biotinylation site by BirA ligase. Four different VP4 proteins tagged with BAP (VP4-BAP) were built between amino acid regions T96-R101 (blue), E109-S114(orange), N132-Q137 (pink), and K145-G150 (green). VP4 trimer ribbon structure for visualization of VP5CT (body and stalk, red) and VP8* (yellow) fragments. An inset in VP8* indicates the different positions in hydrophobic loops of VP8* where the inserted BAP tags were colored in blue, orange, pink, and green. **(B)** Western blot retardation assay of cell lysates transiently expressing wtVP4 and VP4-BAP tagged at blue, orange, pink, and green positions, respectively. Untreated (-) and treated (+) samples with streptavidin are indicated. Immunoblot was incubated with anti-VP4 to detect unbound and bound VP4 to streptavidin (VP4-BAP•StAv). Alpha-tubulin was used as a loading control. **(C)** Comparison of the dsRNA genome segments migration pattern of rRV/wt and rRV/VP4-BAP. The red arrow points to gs -BAP. **(D)** Virus replication fitness curve between 0 to 48 hpi of rRV/wt and rRV/VP4-BAP. **(E)** Immunoblotting of uninfected (-) or infected cell lysates in MA-cytBirA (left panel) or MA104 (right panel) with rRV/wt or rRV/VP4-BAP [MOI, 25 VFU/cell]. Biotinylated proteins were detected with StAv-HRP. Alpha-tubulin was used as a loading control. The red star and red arrow indicate biotinylated VP4-BAP and host undetermined biotinylated protein, respectively. **(F)** WB-ra of MA-cytBirA cell lysates infected with rRV/wt or rRV/VP4-BAP untreated or treated with StAv. The membrane was incubated with anti-VP4 and anti-VP6 for the detection of virus proteins. Alpha-tubulin was used as a loading control. The percentage of biotinylated VP4 normalized according to VP6 expression is indicated. **(G)** WB-ra of purified rRV/VP4-BAP particles incubated without (-) and with (+) streptavidin. The membrane was incubated for the detection of VP4-BAP (anti-VP4) and VP6 (anti-VP6). Alpha-tubulin was used as a loading control. The percentage of biotinylated VP4-BAP was determined and normalized to the expression of VP6. **(H)** Visualization at a high resolution of purified virions isolated of rRV/VP4-BAP infected MA-cytBirA cells untreated (-biotin, upper panel) or treated (+biotin, lower panel) with 100μM biotin. After purification, the virions were labeled with streptavidin conjugated to colloidal gold (12 nm), followed by negative staining and visualization at the electron microscope (right panel). Scale bar is 100 nm

We next assessed whether these four VP4-BAP proteins could assemble into infectious rotavirus particles and support virus replication. For this purpose, we took advantage of a newly developed reverse genetics system to rescue recombinant rotavirus (rRV) harboring a genetically modified genome segment 4 (gs4) encoding the different VP4-BAP proteins (gs4-BAP) (38–41). We were able to rescue only the rRV harboring gs4-BAP encoding the BAP tag within the “green” loop (**Fig. 1A and C**), herein named rRV/VP4-BAP, as demonstrated by the larger size of the modified gs4-BAP compared to the wt gs4 in the dsRNA virus genome migration pattern (**Fig. 1C**) and confirmed by Sanger sequencing (**Fig. S1C**). This outcome suggests that the “green” loop is the only one that preserves virus infectivity when modified by the BAP tag. The rRV/VP4-BAP replication kinetic was delayed compared to the recombinant wt strain (rRV/wt) (**Fig. 1D**), even though similar viral titers were reached at 48 hours post-infection (hpi), suggesting that the VP4-BAP integration in the virus particle affects the virus assembly rate.

We then investigated the ability of VP4-BAP produced by rRV/VP4-BAP to be biotinylated in cells expressing the BirA enzyme. For this purpose, we generated MA104 cells stably expressing cytosolic localized BirA (MA-cytBirA) and infected them with rRV/VP4-BAP. The produced VP4-BAP protein showed biotinylation as demonstrated by a band of approx. 85 kDa detected after incubation with StAv-peroxidase (**Fig.1E, lane 3**). As expected, VP4 biotinylation was detected neither in rRV/wt infected MA-cytBirA cells (**Fig. 1E, lane 2**) nor in rRV/VP4-BAP infected MA104 cells (**Fig. 1E, lanes 4, 5**). Using the WB-ra, we found that the fraction of biotinylated VP4-BAP corresponded to 73% of the total protein (**Fig. 1F**).

We next examined if biotinylated VP4-BAP was able to be incorporated into newly assembled virus particles. We therefore purified virions produced in MA-cytBirA cells in the presence of biotin and estimated their biotinylation by the WB-ra. Consistent with our observation in cell extracts, 67% of the VP4-BAP in virus particles was biotinylated (**Fig. 1G**). We additionally visualized biotinylated VP4-BAP on purified virions by negative staining electron microscopy followed by incubation with StAv conjugated to gold particles. Thus, the virions produced in the presence of biotin were positive to the gold particles (53 %) (**Fig. 1H**). Interestingly, we also identified the presence of virus coat-like layers by negative staining of purified rRV/VP4-BAP particles (**Fig. S1D**) that were absent in samples of the purified rRV/wt, suggesting instability of the rRV/VP4-BAP virions. Moreover, as shown in **Fig. S1E**, these particles appear to have a slightly larger diameter (~80 nm) when compared to rRV/wt particles (~75 nm) but were still in the range of TLPs (42).

### Internalization of rRV/VP4-BAP and cytosolic localization of VP4-BAP

Since the virus replication fitness of rRV/VP4-BAP was delayed compared to rRV-wt, we investigated whether this was caused by a difference in virion internalization. Purified rRV/VP4-BAP and rRV/wt virions labeled with StAv-Alexa 555 before infection were compared and analyzed for virus particle internalization by CLSM. As a control, the virus particles were also immunostained with the conformational monoclonal antibody (mAb) anti-VP7 (clone 159), which only recognizes the trimeric form of the VP7 protein (43, 44). Initially (0 min), VP4-BAP and VP7 signals co-localized on the cell surface, indicating association of virions to the cell membrane, while after two minutes at 37°C, both signals were found already internalized (**Fig. 2A**). These localization patterns were comparable to the ones observed for the same time points with rRV/wt virions (anti-VP7, clone 159), suggesting no differences in the internalization mechanism between both viruses.

**Figure 2.**
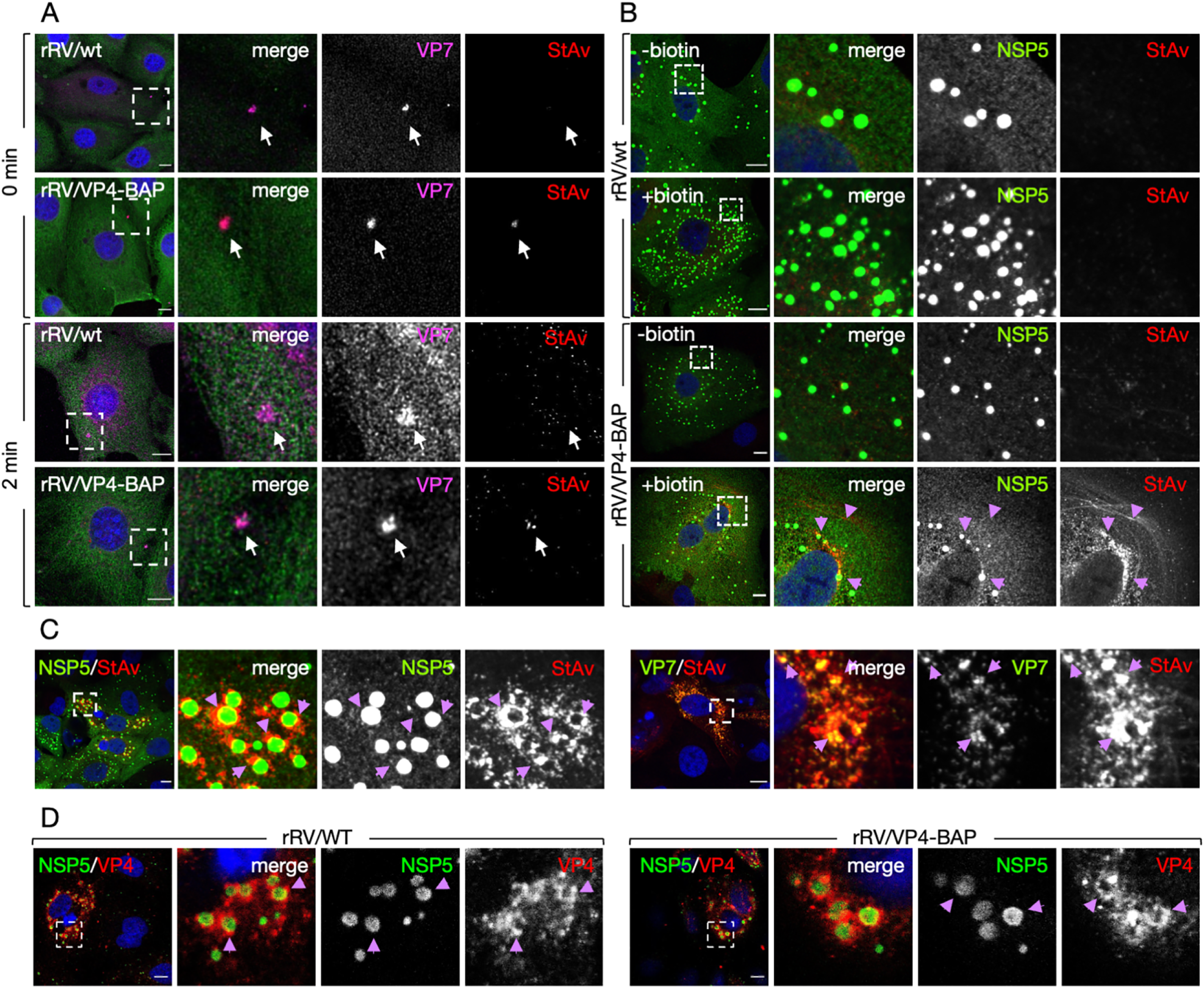
virus entry and RV protein localization upon rRV/VP4-BAP infection. **(A)** Internalization in MA104 cells of purified virions at 0 min (upper panel) and 2 min (lower panel). Purified virions of rRV/wt and biotinylated rRV/VP4-BAP were previously labeled with StAv-Alexa 555 (red). At the indicated time points, cells were fixed and immunostained for VP7 trimers detection (mAb anti-VP7 clone 159, pink) and MTs (anti-α-tubulin, green). Nuclei were stained with DAPI (blue). White open boxes indicate the magnified images at the right. Arrows point to virus particle clamps detected with VP7. Scale bar is 20 μm. **(B)** MA-cytBirA cells infected with rRV/wt (top panel) and rRV/VP4-BAP (bottom panel) untreated (-biotin) and treated (+biotin) with biotin. Cells were PFA fixed at 6 hpi and stained for viroplasms (anti-NSP5, green) and biotinylated proteins (streptavidin-Alexa 555, red) detection. Nuclei were stained with DAPI (blue). **(C)** Immunostaining images of rRV/VP4-BAP infected MA-cytBirA cells in the presence of biotin. At 6 hpi, PFA fixed cells were stained for the detection of VP4-BAP (StAv, red) with viroplasms (anti-NSP5, green) (left row) or mature RV particles (anti-VP7 clone 159, green) (right row). **(D)** immunofluorescence images comparing localization of VP4 and VP4-BAP (anti-VP4, red) of cells infected with rRV/wt (left row) or rRV/VP4-BAP (right row). Viroplasms were detected with anti-NSP5 (green). The dashed white boxes correspond to the image insets of the right columns. Purple arrows point to the VP4-BAP streptavidin signal. Scale bar is 10 μm.

We then compared the localization of the newly produced biotinylated VP4-BAP in rRV/VP4-BAP infected MA-cytBirA cells at 6 hpi, a time point with well-assembled viroplasms (45). For this purpose, infected cells were incubated with or without biotin for 4 hours before fixation. Biotinylated VP4-BAP, detected with StAv-Alexa 555, was found close to viroplasms (revealed with anti-NSP5) and forming bundle-like structures presumably because of the association of VP4 with microtubules and actin filaments (26, 29) (**Fig. 2B**). As expected, no StAv-Alexa 555 signal was detected in cells infected with rRV/wt or with rRV/VP4-BAP in the absence of biotin. Notably, the biotinylated VP4-BAP was found surrounding viroplasms and co-localizing with VP7 in the endoplasmic reticulum (ER) (**Fig. 2C-D**), suggesting that the modification exerted in VP4-BAP does not impact VP4 subcellular localization during RV replication.

### rRV/VP4-BAP revealed a role of VP4 in viroplasm stability

We noticed a different behavior of viroplasms formed by rRV/VP4-BAP or rRV/wt upon fixation of infected cells with methanol. More precisely, few intact viroplasms were detectable in cells infected with rRV/VP4-BAP in contrast to rRV/wt viroplasms, which remained as globular cytosolic inclusions (**Fig. S2A**). Paraformaldehyde fixation did not show differences between the two viruses suggesting a susceptibility of rRV/VP4-BAP viroplasms to alcohols. Since viroplasms have properties of liquid-liquid phase-separated (LLPS) condensates (45–47), and rRV/VP4-BAP viroplasms are less stable, we hypothesized that VP4 may have a yet unidentified role in the viroplasm stability. To challenge this hypothesis, we used 1,6-hexanediol (1,6-HD), a well-described aliphatic alcohol able to disrupt weak hydrophobic protein-protein or protein-RNA interactions, which are key drivers of liquid-liquid phase separation (48, 49) and recently shown to be effective in dissolving RV viroplasms (50). To visualize viroplasms formation in living cells, we took advantage of our previously established MA104 cell line stably expressing NSP2 fused to the monomeric fluorescent protein mCherry (herein named MA-NSP2-mCherry), which is recruited into viroplasms during RV infection (40, 45, 46). Upon infection of MA-NSP2-mCherry cells with either rRV/VP4-BAP or rRV/wt followed by addition of 1,6-HD for 6 min at 5 hpi (**Fig. 3A**), the viroplasms formed by both rRVs dissolved and then readily recovered by 30 min after the compound was washed out (**Fig. S2B**). Interestingly, the rRV-VP4-BAP viroplasms had a delayed recovery kinetic compared to those from rRV/wt at short times after removing the drug (2 min) as denoted by quantifying either the numbers of cells showing viroplasms (**Fig. 3B**) or the numbers of viroplasms per cell (**Fig. 3C**), despite the reduced numbers of viroplasms present in cells infected with rRV-VP4-BAP (**Fig. 3D**) just before the addition of 1,6 HD. Also, the initial size of rRV/VP4-BAP viroplasms was significantly smaller than that of the rRV/wt viroplasms (**Fig. 3E**) and showed delayed size recovery upon 1,6 HD removal (**Fig. 3F**). Notably, at 2 min post-recovery, the viroplasm perinuclear localization was delayed as well for the virus with the tagged VP4 (**Fig. S2C-D**).

**Figure 3.**
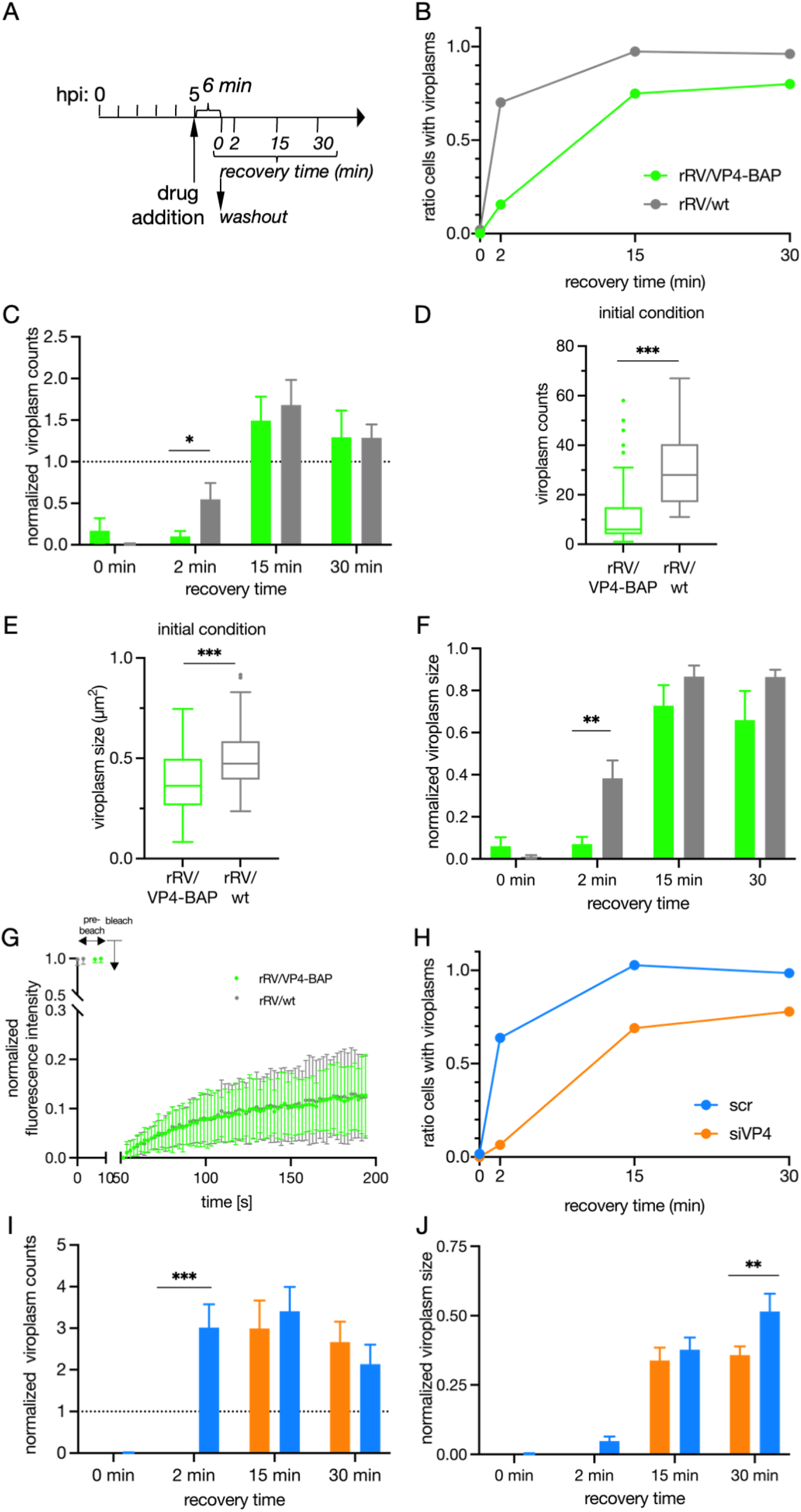
rRV/VP4-BAP viroplasms revealed a delayed dynamic associated with VP4 role. **(A)** Schematic representation for the characterization of LLPS condensates on viroplasms of rRV/VP4-BAP-infected cells. At 5 hpi, RV infected MA/NSP2-mCherry cells were treated with 1,6-HD for 6 min. The drug was washed out, and samples were fixed and imaged for viroplasm quantification at 0-, 2-, 15- and 30-min post-recovery. **(B)** 1,6-HD recovery plot of cells showing viroplasms normalized to initial conditions (5 hpi). **(C)** 1,6-HD recovery plot of viroplasm counts per cell upon infection with rRV/VP4-BAP (green column) and rRV/wt (grey column). Plots of viroplasm counts per cell **(D)** and viroplasm size per cell **(E)** at initial conditions (5 hpi). **(F**) 1,6-HD recovery plot of normalized viroplasm size at initial conditions. **(G)** FRAP recovery curve of NSP2-mCherry of single viroplasms of rRV/VP4-BAP (green) and rRV/wt (grey) infected MA/NSP2-mCherry cells, 5 hpi (n=27 and 25, respectively). 1,6-HD recovery plots showing appearing viroplasms **(H)**, number of viroplasms per cell **(I)**, and viroplasm size per cell **(J)** of SA11 infected MA104 cells silenced with siVP4 (orange) or control siRNA (scr, blue). Data represent mean ± SEM Student’s t-test (*), p<0.05; (**), p<0.01 and (***), p<0.001

Next, we wanted to examine whether the liquid-like properties of rRV/VP4-BAP viroplasms were altered. For this purpose, we measured the NSP2-mCherry diffusion dynamics in single viroplasms using fluorescence recovery after photobleaching (FRAP) experiments (**Fig. 3G and S2E**). Surprisingly, we found that the viroplasms fluorescence recovery after photobleaching at 5 hpi was similar for both viruses. Moreover, no differences were observed in the half-time recovery or mobile fraction. (**Fig. S2F-G**). Thus, these results exclude differences in the liquid-like properties of these globular inclusions. In order to further characterize the relationship between viroplasms and VP4, we used a siRNA depletion approach. Although silencing of VP4 was previously shown to impair assembly of the virion-third layer leading to the accumulation of DLPs, viroplasms were still formed (**Fig. S3A-B**) (51). We thus reasoned that in the absence of VP4, viroplasms should have similar behavior as those observed during rRV/VP4-BAP infection. Although methanol fixation of viroplasms in VP4 silenced cells and infected with RV strain SA11 did not substantially affect viroplasm morphology to the same extend as that of rRV/VP4-BAP (**Fig. S3C**). Specifically, the viroplasms in the cells infected with the wt virus showed a diffuse morphology while those in rRV/VP4-BAP completely dissolved. As described above, experiments performed with 1,6-HD (**Fig. S3D**) had a similar delayed viroplasm recovery kinetic on VP4-less viroplasms as observed for rRV/VP4-BAP viroplasms **(Fig. 3H)**. Moreover, the number (**Fig. 3I**) and the size (**Fig. 3J**) of viroplasms were both decreased in siVP4 treated cells when compared to the experimental controls.

Collectively, our data indicate that VP4 can be modified by the insertion of relatively short tags at least in one position within the VP8* lectin domain of VP4 and suggest that VP4 plays a role in the stability and dynamics of the viroplasms.

## DISCUSSION

The external coat layer of the RV virion can be modified by adding *in vitro* a specific ratio of VP7 and VP4 proteins to purified DLPs to generate recoated TLPs (rcTLPs) (10–12, 52), which is a valuable tool to study the VP4 structural requirements allowing virion internalization (10). However, rcTLPs have methodological limitations and do not allow transferring the parental phenotype to the virus progeny. Moreover, rcTLPs only allow single amino acid substitutions of the spike VP4 (53, 54). Here, we used a RV reverse genetics system (40) to show that it is possible to modify an RV structural protein and specifically to remodel the spike protein VP4. For this purpose, four exposed loops present in the VP8* subunit lectin domain were modified by incorporating a short BAP tag of 15 amino acids. Interestingly, although the four differently modified proteins could be efficiently biotinylated in transfected cells, only one of them, with a BAP tag inserted in the K145-G150 loop, could be incorporated into infectious viral particles and rescued by reverse genetics. The addition of BAP tags in other positions of the VP8* subunit may destabilize the VP4 structure, interfering with its incorporation into the virion and, thus, not allowing the rescue of infectious viruses. We reasoned, therefore, that these VP4-BAP versions are strongly compromise newly generated virions because they might directly impact: *i*) the transition from upright (immature) to reverse (mature) conformational VP4 states (53); *ii*) the association with specific cellular receptors (19, 20) or, *iii*) the incorporation of VP4 in the coat layer.

The internalization of RV virions requires a sequence of events involving interaction with the cell-membrane, followed by invagination, and then engulfment into endosomes. This process triggers a decrease of calcium levels within the endosomes, which induces loss of the virion VP4-VP7 outer layer and the release of transcriptionally active DLPs into the cytosol (54–57). The internalization kinetics of the rRV/VP4-BAP, as denoted by the ability to reach the cytosol, were found to be comparable to that of the rRV/wt. Also, VP4-BAP and VP4 share similar distribution patterns in infected cells, such as localization surrounding viroplasms, co-localization in the endoplasmic reticulum with VP7, and incorporation into newly assembled virions. However, we found that rRV/VP4-BAP has a reduced virus replication fitness compared to rRV/wt. Intriguingly, rRV/VP4-BAP viroplasms seemed to be less resistant to methanol fixation than rRV/wt, which led us to analyze viroplasms in the context of LLPS condensates by using the well-described 1,6-HD (48, 49), which compromises RV viroplasms integrity. Upon 1,6-HD removal, the re-assembly of rRV/VP4-BAP viroplasms was slower, and their number and size were reduced compared to rRV/wt. However, no differences in viroplasm liquid-like dynamics were found when the mobility of NSP2-mCherry was analyzed on single-viroplasm FRAP experiments, suggesting that the reduction observed for rRV/VP4-BAP was related to a flawed process in the viroplasm assembly but not to already formed inclusions. Our results are consistent with viroplasm behavior as LLPS condensates(50). The only difference between rRV/VP4-BAP and the rRV/wt resides in the structure of VP4, implying a role of VP4 in the structural stability of viroplasms. On this line of thinking, the rRV/VP4-BAP viroplasms behavior was further confirmed by silencing VP4 in SA11-infected cells. Consistent with a previous publication (51), the viroplasm formation was not affected by the depletion of VP4. Nevertheless, the effect on viroplasm recovery kinetic after 1,6-HD treatment was similar to the one observed on rRV/VP4-BAP viroplasms. Altogether, our data support a new functional role of VP4 directly linked to the stabilization and assembly of viroplasms. In fact, VP4 has been described to interact with the actin cytoskeleton and the RV restrictive factor drebrin (25, 29, 30). Interestingly, a highly conserved actin-binding domain present in the C-terminus of VP4 has been shown to remodel actin bundles to favor RV exit (27, 28). Most of these VP4-cytoskeleton associations involve the VP5* subunit. Intriguingly, since the alteration of viroplasm stability was observed upon modification of the VP8* subunit, we hypothesize that the VP8* subunit is involved in at least one of these three aspects that render the viroplasms assembled and stabilized: *i*) association of VP8* with a yet undescribed host component, *ii*) a reorganization ofVP5*-VP8* association or *iii*) a direct role of VP8* over another RV protein(s).

Identifying a target site in the spike VP4 (loop region K145-G150) permissive for the insertion of an exogenous peptide may impact the RV field. This VP4 modification favors the insertion of peptides required for super-resolution microscopy or DNA-paint technologies (e.g., Halo or BC2 tags) to dissect debated aspects of RV entry. In addition, this VP4 modification technology could permit the incorporation of antigenic peptides for vaccine development. Although it is well-known that the current oral RV vaccines elicit an immune response (58, 59), the use of rRV harboring a modified VP4 could provide an improved vaccination platform for the display of other antigens fostering the development of a new generation of dual-vaccines.

## MATERIALS AND METHODS

### Cells and viruses

MA104 cells (embryonic African green monkey kidney cells; ATCC CRL-2378) were grown in Dulbecco’s modified Eagle’s medium (DMEM) (Life Technologies) containing 10% fetal calf serum (FCS) (AMIMED; BioConcept, Switzerland) and penicillin (100 U/ml)–streptomycin (100 μg/ml) (Gibco, Life Technologies). MA/cytBirA and MA/NSP2-mCherry (40) cell lines were grown in DMEM supplemented with 10% FCS, penicillin (100 U/ml)-streptomycin (100μg/ml) and 5μg/ml puromycin (InvivoGen, France). BHK-T7/9 (baby hamster kidney stably expressing T7 RNA polymerase) cells were kindly provided by Naoto Ito (Gifu University, Japan)(60) and cultured in Glasgow medium supplemented with 5% FCS, 10% tryptose phosphate broth (Sigma-Aldrich), 10% FCS, penicillin(100 U/ml)-streptomycin (100μg/ml), 2% nonessential amino acids and 1% glutamine.

rRV/wt (40), rRV/VP4-BAP, and simian rotavirus strain SA11 (G3P6[1])(61) were propagated, grown, and purified as previously described (62). Virus titer was determined as viroplasm forming units per ml (VFU/ml) as described by (45). The T_7_ RNA polymerase recombinant vaccinia virus (strain vvT7.3) was amplified as previously described (37).

### Cell line generation

MA/cyt-BirA cell line was generated using the PiggyBac technology (63). Briefly, 10^5^ MA104 cells were transfected with the pCMV-HyPBase (63) and transposon plasmids pPB-cytBirA using a ratio of 1:2.5 with Lipofectamine 3000 (Sigma-Aldrich) according to the manufacturer’s instructions. The cells were maintained in DMEM supplemented with 10% FCS for three days and then incubated with DMEM supplemented with 10% FCS and 5 μg/ml puromycin (Sigma-Aldrich) for four days to allow the selection of cells expressing the gene of interest (40).

### Reverse genetics

rRV/VP4-BAP was prepared as described previously (40, 41) using a pT_7_-VP4-BAP instead of pT_7_-VP4. Briefly, monolayers of BHK-T_7_ cells (4 × 10^5^) cultured in 12-well plates were co-transfected using 2.5 μL of TransIT-LT1 transfection reagent (Mirus) per microgram of DNA plasmid. The mixture comprised 0.8 μg of SA11 rescue plasmids: pT_7_-VP1, pT_7_-VP2, pT_7_-VP3, pT_7_-VP4-BAP, pT_7_-VP6, pT_7_-VP7, pT_7_-NSP1, pT_7_-NSP3, pT_7_-NSP4, and 2.4 μg of pT_7_-NSP2 and pT_7_-NSP5 (38, 39). Additionally, 0.8 μg of pcDNA3-NSP2 and 0.8 μg of pcDNA3-NSP5, encoding NSP2 and NSP5 proteins, were co-transfected to increase rescue efficiency (40, 41). Cells were co-cultured with MA104 cells for three days in serum-free DMEM supplemented with trypsin from porcine pancreas (0.5 μg/ml final concentration) (T0303-Sigma Aldrich) and lysed by freeze-thawing. 300 μL of the lysate was transferred to fresh MA104 cells and cultured at 37°C for four days in serum-free DMEM supplemented with 0.5 μg/ml trypsin until a visible cytopathic effect. The modified genome segments of rescued recombinant rotaviruses were confirmed by specific PCR segment amplification followed by sequencing (40).

### Antibodies and Chemicals

Guinea pig anti-NSP5, guinea pig anti-RV, and rabbit anti-VP4 were described previously (45, 64, 65). Rabbit anti-NSP3 was kindly provided by Susana Lopez (UNAM, Mexico), Mouse monoclonal anti-VP7 (clone 159) was kindly provided by Harry Greenberg (Stanford University, CA, USA). Mouse mAb anti-glyceraldehyde dehydrogenase (GAPDH) (clone GAPDH-71.1) and mouse anti-alpha tubulin (clone B-5-1-12) were purchase to Merck. Streptavidin-HRP was purchased from Merck. Streptavidin-Alexa 555 and secondary antibodies conjugated to Alexa 488, Alexa 594, Alexa 647, Alexa 700 (ThermoFisher Scientific).

### DNA plasmids

pcDNA-VP4-SA11 was obtained by RT-PCR amplification of VP4 ORF of gs 4 from rotavirus simian strain SA11 (66) using specific primers to insert *Hind*III and *Xho*I sites, followed by ligation into those sites in pcDNA3 (Invitrogen). pcDNA-VP4-*Kpn*I/*BamH*I was built by insertion of point mutations in pcDNA-VP4-SA11 using the QuikChange site-directed mutagenesis kit and protocol (Agilent) to insert *Kpn*I and *BamH*I restriction sites in VP4. pcDNA-VP4-BAP (blue), (orange), (pink), and (green) were obtained by ligation between *Kpn*I and *BamH*I of pcDNA-VP4-*Kpn*I/*BamH*I a synthetic DNA fragment (GenScript®) containing BAP tag in VP4 loops in amino acid regions 96-101, 109-114, 132-137 and 145-150, respectively. The BAP tags are flanked by *BspE*I and *Nhe*I restriction sites for easy tag replacement. A detailed list of used DNA sequence fragments is in ***SI Appendix, Table 1***.

RV plasmids pT_7_-VP1-SA11, pT_7_-VP2-SA11, pT_7_-VP3-SA11, pT_7_-VP4-SA11, pT_7_-VP6-SA11, pT_7_-VP7-SA11, pT_7_-NSP1-SA11, pT_7_-NSP2-SA11, pT_7_-NSP3-SA11, pT_7_-NSP4-SA11, and pT_7_-NSP5-SA11 were previously described (38). pcDNA3-NSP5 and pcDNA3-NSP2 were already described (40). pT_7_-VP4-BAP (blue), (orange), (green), and (pink) were obtained by inserting a synthetic DNA fragment (Genscript) encoding for the VP4 protein-encoding BAP tag flanked by *Mfe*I and *Nde*I restriction enzymes sites and ligated into those sites in the pT_7_-VP4-SA11. A list of the synthesized DNA fragment is *SI Appendix, Table 1*.

pPB-cytBirA plasmid was obtained from a synthetic DNA fragment (Genscript) containing the BirA enzyme open reading frame of *Escherichia coli* (UniProt accession number: P06709) and inserted in the pPB-MCS vector (41) using *Nhe*I-*BamH*I restriction enzymes sites.

### Streptavidin-supershift assay

The assay was performed as described by Predonzani et al.(34). Briefly, cell extracts were lysed in TNN lysis buffer (100mM Tri-HCl pH8.0, 250 mM NaCl, 0.5% NP-40, and cOmplete protease inhibitor (Roche)) and centrifuged for 7 min at 15’000 rpm and 4°C. The supernatant was exhaustively dialyzed against PBS (phosphate-buffered saline, 137 mM NaCl, 2.7 mM KCl, 8 mM Na_2_HPO_4_, and 2 mM KH_2_PO_4_ pH 7.2) at 4°C and heated for 5 min at 95°C in Laemmli sample buffer. Samples were incubated for 1 h at 4°C with 1 μg streptavidin (Sigma) and then resolved in SDS-polyacrylamide gel under reducing conditions. Proteins were transferred to nitrocellulose 0.45 μm (67) and incubated with corresponding primary and secondary antibodies. Secondary antibodies were conjugated to IRDye680RD or IRDye800RD (LI-COR, Germany) for protein detection and quantification in Odyssey® Fc (LI-COR Biosciences).

### Virus fitness curve

The experiment was performed as described previously (47) with some modifications. MA104 cells (2 × 10^5^) seeded in 12-well plates were infected with rRV at an MOI of 10 VFU/cell. The virus was allowed to adsorb for 1 h at 4°C, followed by incubation at 37°C in 500 μl DMEM. At the indicated time points, the plates were frozen at −80°C. The cells were then treated with three freeze-thaw cycles, harvested, and centrifuged at 17,000 × *g* for 5 min at 4°C. The supernatant was recovered and activated with 80 μg/ml of trypsin for 30 min at 37°C. Two-fold serial dilutions were prepared and used to determine the viral titers described previously (40, 41).

### Fluorescence labeling of purified rRV

100 μl of purified biotinylated rRV/VP4-BAP is activated for 30 min at 37°C with 4 ul trypsin (2 mg/ml). The mixture is then incubated with 1 μl of streptavidin-Alexa Fluor 555 (2mg/ml) (ThermoFisher Scientific) for 1 h at room temperature. The tube was snaped every 20 min. Unbound streptavidin was separated labeled virus by loading the 50 μl reaction mixture on top of 100 μl of a 20% sucrose-PBS cushion. Samples were centrifuged for 40 min at 20 psi on Airfuge air-driven ultracentrifuge (Beckman Coulter). Pellet was resuspended in 20 μl Tris-buffered saline (TBS) buffer (25 mM Tris-HCl, pH 7.4, 137 mM NaCl, 5 mM KCl, 1 mM MgCl_2_, 0.7 mM CaCl_2_, 0.7 mM Na_2_HPO_4_, 5.5 mM dextrose).

### Immunofluorescence

For virus internalization experiments, 1 μl of rRV particles conjugated to SA-Alexa555 diluted in 50 μl of DMEM was adsorbed over MA104 cells for 15 min in a metal tray cooled to −20°C. Cells were then transferred to 37°C and fixed at the indicated time-points with ice-cold methanol for 3 min on dry ice.

For later times post-infection, the virus was adsorbed for 1h at 4°C in a reduced volume. Then, cells were transferred to 37°C, treated at the indicated time points with 100μM biotin in DMEM serum-free. Cells were fixed in 2% paraformaldehyde in phosphate-buffered saline (PBS) for 10 min at room temperature. All immunofluorescences were processed as described by Buttafuoco *et al*. (67). Images were acquired using a confocal laser scanning microscope (CLSM) (DM550Q; Leica). Data were analyzed with the Leica Application Suite (Mannheim, Germany) and Image J (68).

### LLPS characterization

MA/NSP2-mCherry cells were seeded at a density of 1.2×10^4^ cells per well 8-wells Lab-Tek® Chamber Slide™ (Nunc, Inc. Cat #177402). For RV infection, the virus was adsorbed at MOI of 25 VFU/cell diluted in 30 μl of DMEM serum-free, incubated at 4°C for 1 h in an orbital shaker and then volume filled to 100μl with DMEM-serum-free followed by incubation at 37°C. At 5 hpi, the media was replaced by media containing 3.5% 1,6-hexanediol (Sigma-Aldrich) in 2% FCS-DMEM and cells were incubated for 6 min at 37°C. Then the drug was washed out by removing the media, washing the cells three times with PBS, and adding fresh 2%FCS-DMEM and incubated at 37°C. At designated time post-recovery, cells were fixed with 2% PFA for 10 min at room temperature. Nuclei were stained by incubating cells with 1 μg/ml of DAPI (4’,6-diamidino-2-phenylindole) in PBS for 15 min at room temperature. Samples were mounted in ProLong™ Gold antifade mountant (Thermo Fischer Scientific), and Images were acquired using a fluorescence microscope (DMI6000B, Leica). Data were analyzed with ImageJ (version 2.1.0/1.53; https://imagej.net/Fiji).

### Quantification of viroplasms

Number, size, and perinuclear localization of viroplasms were essentially acquired and analyzed as previously described (45, 69–71). The viroplasm perinuclear ratio was determined as previously described (69, 70) using the following formula: (V-N)/N, whereas V, area occupied by viroplasm and N, area of the nucleus. Data analysis was performed using Microsoft® Excel (version 16.46), and the statistical significance of differences was determined by unpaired parametric Welch’s t-test comparison post-test, using Prism 9 (GraphPad Software, LLC).

### Rotavirus genome pattern visualization

Rotavirus genome extraction and visualization were performed as previously described (40, 41).

### Negative staining of purified particles

For staining of biotinylated TLPs with streptavidin-gold, purified particles were dialyzed overnight at 4°C in TNC buffer (10 mM Tris-HCl, pH 7.5, 140 mM NaCl, 10mM CaCl2). The TLPs were adsorbed for 10 min on carbon-coated Parlodion films mounted on 300-mesh copper grids (EMS). Samples were washed once with water, fixed in 2.5% glutaraldehyde in 100 mM Na/K-phosphate buffer, pH 7.0, for 10 min at room temperature, and washed twice with PBS before incubation with 10 μl streptavidin conjugated to 10 nm colloidal gold (Sigma-Aldrich, Inc) for 2 h at room temperature. The streptavidin-gold conjugated was treated as described previously (63) before use to separate unconjugated streptavidin from streptavidin-conjugated (72) to colloidal gold. The viral particles were further washed three times with water and stained with 2% phosphotungstate, pH 7.0 for 1 min at room temperature. Samples were analyzed in a transmission electron microscope (CM12; Philips, Eindhoven, The Netherlands) equipped with coupled device (CCD) cameras (Ultrascan 1000 and Orius SC1000A; Gatan, Pleasanton, CA, USA) at an acceleration voltage of 100 kV.

For calculation of the diameter of virus particles by negative staining, the area of each virus particle was calculated using Imaris software (version 2.1.0/1.53c; Creative Commons license) and then converted to the diameter as follow: π, where a is the area and d is the diameter of the particle, respectively.

### siRNA reverse transfection

For silencing gs 4 of SA11strain, the following siRNA pool: siVP4-25 (5’-UUGCUCACGAAUUCUUAUATT-3’), siVP4-931(5’-GAAGUUACCGCACAUACUATT-3’) and siVP4-1534 (5’-AUUGCAAUGUCGCAGUUAATT −3’ pool was designed and synthesized by Microsynth AG (Switzerland). siRNA-A (sc-37007, Santa Cruz Biotechnology) was used as scrambled siRNA. siRNA reverse transfection was performed by mixing 1.2 μl siRNA 5 μM with 1 μl lipofectamine RNAiMAX transfection reagent (Invitrogen, ThermoFisher Scientific) to a final volume of 100 μl with Opti-MEM® (Gibco, ThermoFisher Scientific) in a well of 24-well plate and incubated for 20 min at room temperature. To reach a 10nM siRNA final concentration, 2×10^4^ cells diluted in 500 μl DMEM supplemented with 10% FCS are added on top and incubated for 60 h previous to analysis. Thus, cells were infected with RV strain SA11 at MOI 12 VFU/cell as described previously (45, 47, 67).

### FRAP

1.2 ×10^4^ MA/NSP2 cells per well were seeded in μ-Slide 18-well glass-bottom plates (Ibidi). Cells were RV-infected at MOI of 15 VFU/cell and kept in DMEM-SF. At 4.5 hpi, the cells were counterstained with Hoechst 33342 diluted in FluoroBRITE DMEM (Gibco, Cat.No. A18967-01) at a concentration of 1μg/ml, incubated for 30 min at 37°C and subjected to FRAP analysis. FRAP experiments were performed with an SP8 Falcon confocal laser scanning microscope (CLSM) from Leica equipped with a 63x objective (NA 1.4) using the FRAP function of the LasX software (Leica) as follows: a circular area of 2 μm in diameter, encompassing an entire viroplasm, was bleached with the 405 nm and 481 nm lasers at 100% laser power for 20 iterations. The fluorescent recovery was monitored by taking fluorescence images of the mCherry channel every 2 seconds for 140 min. For each FRAP acquisition, a circular area of 2 μm, encompassing an entire unbleached viroplasm in the same cell, was used as the fluorescent control, and a squared area of 5 μm x 5 μm was chosen as background. The entire FRAP dataset was analyzed with MatLab (MATLAB R2020b, Mathworks) using the FRAP-tool source code from easyFRAP (Cell Cycle Lab, Medical School, University of Patras). Fully normalized data were used to generate FRAP diagrams and calculate recovery half-times (T-half) and mobile fractions from independent measurements. Representative images were taken and processed for each FRAP experiment using the Imaris software v9.5 (Bitplane, Oxford Instruments). Fluorescent intensities of FRAP movies were normalized using a customized Fiji pipeline (68).

## ACKNOWLEDGMENTS

This work has been supported by the University of Zurich. This project was also supported by a pre-doctoral ICGEB fellowship to GP and GDL. KG was supported by RAV Zollikofen and Diaconis - AMM Berner Stellennetz, Switzerland.

The authors declare no conflict of interest.

## SUPPLEMENTAL MATERIAL

### SUPPLEMENTARY FIGURE LEGENDS

**Figure S1. (A)** Immunoblotting of cellular extracts of transfected VP4-BAP (lane 1) and infected rRV/VP4-BAP (lane 2). **(B)** Immunoblot of cellular extracts of transfected VP4-BAP untreated (-) or treated (+) with lambda phosphatase. VP4-BAP shifted band is indicated with a red start. Membranes were incubated with anti-VP4 and anti-tubulin. **(C)** Sequence chromatogram of gs4-BAP of rRV/VP4-BAP visualizing inserted linkers and BAP tag. Nucleotide (top) and amino acid (bottom) sequences are indicated. **(D)** Negative staining of virion layers detected in purified rRV/VP4-BAP preparations. The yellow arrow points to TLPs. Scale bar is 100 nm. **(E)** Scatter dot plot comparing the diameter of purified particles from non-biotinylated (-biotin, light green) or biotinylated (+biotin, dark green) RV/VP4-BAP and rRV/wt (grey). The median value is indicated, n>40 particles, t-test student, (*) p-value<0.05.

**Figure S2. (A)** Immunofluorescence images of viroplasms (anti-NSP5, green) from cells infected with rRV/wt (left panel) or rRV/VP4-BAP (right panel). At 6 hpi, cells were fixed with either PFA (upper row) or methanol (lower row), followed by immunostaining. Nuclei were stained with DAPI (blue). White arrowheads point to green aggregates. Scale bar is 20 μm. **(B)** Representative images of MA-NSP2-mCherry cells infected at 5 hpi with rRV/wt (upper row) or rRV/VP4-BAP (lower row) and treated for 6 min with 3.5% of 1,6-HD. Cells were washed and monitored for viroplasm formation at 0-, 2-, 15- and 30-min post-recovery. White arrows point to cells showing recovered viroplasms. Scale bar is 10 μm. **(C)** Plot for the perinuclear ratio of cells infected with rRV/VP4-BAP and rRV/wt at initial conditions (5 hpi). **(D)** 1,6-HD recovery plot of the normalized perinuclear ratio of viroplasms from cells infected with rRV/VP4-BAP (green) and rRV/wt (grey). **(E)** Fluorescence images of FRAP measurement of single viroplasms of cells infected with rRV/VP4 (top) and rRV/wt (bottom) at pre-bleach, post-bleach, and recovery time conditions. Each inset indicates the bleached viroplasm of the images at the right. Nuclei were stained with Hoescht 33342. Scale bar is 10 μm. Plots of the T-half recovery **(F)** and the mobile fraction **(G)** means of single viroplasms of rRV/VP4-BAP and rRV/wt.

**Figure S3. (A)** Immunoblot of 6 hpi cellular lysates prepared from MA104 or MA-NSP2-mCherry cells silenced with siVP4 or control siRNA (scr) followed by mock-infection or infection with RV simian strain SA11. The membrane was stained with anti-VP4, anti-NSP5, and anti-GAPDH (loading control). **(B)** Immunostaining at 6 hpi of SA11-infected MA-NSP2-mCherry cells knocked down with control siRNA (scr) (upper row) or siVP4 (lower row). Cells were immunostained with anti-VP4 (green). Nuclei were stained with DAPI (blue). Scale bar is 10 μm. **(C)** Immunofluorescence analysis at 6 hpi of SA11-infected MA104 cells silenced with siVP4 or control siRNA. Cells were fixed either PFA (upper row) or methanol (lower row), followed by viroplasm immunostaining (anti-NSP5, green). Nuclei were stained with DAPI (blue). White arrowheads point to diffuse viroplasms. Scale bar is 20 μm. **(D)** Representative images of SA11-infected MA-NSP2-mCherry cells knocked down with scr (upper row) or siVP4 (lower row) and treated for 6 min with 3.5% of 1,6-HD. Cells were washed and monitored for viroplasm formation at 0-, 2-, 15- and 30-min post-recovery. White arrows point to cells showing recovered viroplasms. Scale bar is 10 μm.

## REFERENCES

1. Troeger C, Khalil IA, Rao PC, Cao S, Blacker BF, Ahmed T, Armah G, Bines JE, Brewer TG, Colombara DV, Kang G, Kirkpatrick BD, Kirkwood CD, Mwenda JM, Parashar UD, Petri WA, Riddle MS, Steele AD, Thompson RL, Walson JL, Sanders JW, Mokdad AH, Murray CJL, Hay SI, Reiner RC. 2018. Rotavirus vaccination and the global burden of rotavirus diarrhea among children younger than 5 years. JAMA Pediatr 172:958–965.

2. Gomez DE, Weese JS. 2017. Viral enteritis in calves. Can Vet J 58:1267–1274.

3. Vlasova AN, Amimo JO, Saif LJ. 2017. Porcine rotaviruses: epidemiology, immune responses and control strategies. Viruses 9: 48.

4. Dhama K, Saminathan M, Karthik K, Tiwari R, Shabbir MZ, Kumar N, Malik YS, Singh RK. 2015. Avian rotavirus enteritis - an updated review. Vet Q 35:142–58.

5. Lawton JA, Zeng CQ, Mukherjee SK, Cohen J, Estes MK, Prasad BV. 1997. Three-dimensional structural analysis of recombinant rotavirus-like particles with intact and amino-terminal-deleted VP2: implications for the architecture of the VP2 capsid layer. J Virol 71:7353–60.

6. Zhang X, Settembre E, Xu C, Dormitzer PR, Bellamy R, Harrison SC, Grigorieff N. 2008. Near-atomic resolution using electron cryomicroscopy and single-particle reconstruction. Proc Natl Acad Sci U S A 105:1867–72.

7. Charpilienne A, Lepault J, Rey F, Cohen J. 2002. Identification of rotavirus VP6 residues located at the interface with VP2 that are essential for capsid assembly and transcriptase activity. J Virol 76:7822–31.

8. Lepault J, Petitpas I, Erk I, Navaza J, Bigot D, Dona M, Vachette P, Cohen J, Rey FA. 2001. Structural polymorphism of the major capsid protein of rotavirus. EMBO J 20:1498–507.

9. Li Z, Baker ML, Jiang W, Estes MK, Prasad BV. 2009. Rotavirus architecture at subnanometer resolution. J Virol 83:1754–66.

10. Settembre EC, Chen JZ, Dormitzer PR, Grigorieff N, Harrison SC. 2011. Atomic model of an infectious rotavirus particle. EMBO J 30:408–16.

11. Yoder JD, Dormitzer PR. 2006. Alternative intermolecular contacts underlie the rotavirus VP5* two- to three-fold rearrangement. EMBO J 25:1559–68.

12. Yoder JD, Trask SD, Vo TP, Binka M, Feng N, Harrison SC, Greenberg HB, Dormitzer PR. 2009. VP5* rearranges when rotavirus uncoats. J Virol 83:11372–7.

13. Trask SD, McDonald SM, Patton JT. 2012. Structural insights into the coupling of virion assembly and rotavirus replication. Nat Rev Microbiol 10:165–77.

14. Arias CF, Romero P, Alvarez V, López S. 1996. Trypsin activation pathway of rotavirus infectivity. J Virol 70:5832–9.

15. Gilbert JM, Greenberg HB. 1998. Cleavage of rhesus rotavirus VP4 after arginine 247 is essential for rotavirus-like particle-induced fusion from without. J Virol 72:5323–7.

16. Dormitzer PR, Nason EB, Prasad BV, Harrison SC. 2004. Structural rearrangements in the membrane penetration protein of a non-enveloped virus. Nature 430:1053–8.

17. Dormitzer PR, Sun ZY, Wagner G, Harrison SC. 2002. The rhesus rotavirus VP4 sialic acid binding domain has a galectin fold with a novel carbohydrate binding site. EMBO J 21:885–97.

18. Fiore L, Greenberg HB, Mackow ER. 1991. The VP8 fragment of VP4 is the rhesus rotavirus hemagglutinin. Virology 181:553–63.

19. Graham KL, Halasz P, Tan Y, Hewish MJ, Takada Y, Mackow ER, Robinson MK, Coulson BS. 2003. Integrin-using rotaviruses bind alpha2beta1 integrin alpha2 I domain via VP4 DGE sequence and recognize alphaXbeta2 and alphaVbeta3 by using VP7 during cell entry. J Virol 77:9969–78.

20. Graham KL, Takada Y, Coulson BS. 2006. Rotavirus spike protein VP5* binds alpha2beta1 integrin on the cell surface and competes with virus for cell binding and infectivity. J Gen Virol 87:1275–1283.

21. Estes MK, Graham DY, Gerba CP, Smith EM. 1979. Simian rotavirus SA11 replication in cell cultures. J Virol 31:810–5.

22. Enouf V, Chwetzoff S, Trugnan G, Cohen J. 2003. Interactions of rotavirus VP4 spike protein with the endosomal protein Rab5 and the prenylated Rab acceptor PRA1. J Virol 77:7041–7.

23. Zárate S, Cuadras MA, Espinosa R, Romero P, Juárez KO, Camacho-Nuez M, Arias CF, López S. 2003. Interaction of rotaviruses with Hsc70 during cell entry is mediated by VP5. J Virol 77:7254–60.

24. Guerrero CA, Bouyssounade D, Zárate S, Isa P, López T, Espinosa R, Romero P, Méndez E, López S, Arias CF. 2002. Heat shock cognate protein 70 is involved in rotavirus cell entry. J Virol 76:4096–102.

25. Li B, Ding S, Feng N, Mooney N, Ooi YS, Ren L, Diep J, Kelly MR, Yasukawa LL, Patton JT, Yamazaki H, Shirao T, Jackson PK, Greenberg HB. 2017. Drebrin restricts rotavirus entry by inhibiting dynamin-mediated endocytosis. Proc Natl Acad Sci U S A 114:E3642–E3651.

26. Nejmeddine M, Trugnan G, Sapin C, Kohli E, Svensson L, Lopez S, Cohen J. 2000. Rotavirus spike protein VP4 is present at the plasma membrane and is associated with microtubules in infected cells. J Virol 74:3313–20.

27. Trejo-Cerro Ó, Eichwald C, Schraner EM, Silva-Ayala D, López S, Arias CF. 2018. Actin-dependent nonlytic rotavirus exit and infectious virus morphogenetic pathway in nonpolarized Cells. J Virol 92:e02076–17.

28. Condemine W, Eguether T, Couroussé N, Etchebest C, Gardet A, Trugnan G, Chwetzoff S. 2019. The C terminus of rotavirus VP4 protein contains an actin binding domain which requires cooperation with the coiled-coil domain for actin remodeling. J Virol 93:e01598–18.

29. Gardet A, Breton M, Fontanges P, Trugnan G, Chwetzoff S. 2006. Rotavirus spike protein VP4 binds to and remodels actin bundles of the epithelial brush border into actin bodies. J Virol 80:3947–56.

30. Gardet A, Breton M, Trugnan G, Chwetzoff S. 2007. Role for actin in the polarized release of rotavirus. J Virol 81:4892–4.

31. Wolf M, Vo PT, Greenberg HB. 2011. Rhesus rotavirus entry into a polarized epithelium is endocytosis dependent and involves sequential VP4 conformational changes. J Virol 85:2492–503.

32. Beckett D, Kovaleva E, Schatz PJ. 1999. A minimal peptide substrate in biotin holoenzyme synthetase-catalyzed biotinylation. Protein Sci 8:921–9.

33. De Lorenzo G, Eichwald C, Schraner EM, Nicolin V, Bortul R, Mano M, Burrone OR, Arnoldi F. 2012. Production of *in vivo*-biotinylated rotavirus particles. J Gen Virol 93:1474–82.

34. Predonzani A, Arnoldi F, López-Requena A, Burrone OR. 2008. *In vivo* site-specific biotinylation of proteins within the secretory pathway using a single vector system. BMC Biotechnol 8:41.

35. Fairhead M, Howarth M. 2015. Site-specific biotinylation of purified proteins using BirA. Methods Mol Biol 1266:171–84.

36. Roux KJ, Kim DI, Burke B. 2013. BioID: a screen for protein-protein interactions. Curr Protoc Protein Sci 74:19.23.1–19.23.14.

37. Fuerst TR, Niles EG, Studier FW, Moss B. 1986. Eukaryotic transient-expression system based on recombinant vaccinia virus that synthesizes bacteriophage T7 RNA polymerase. Proc Natl Acad Sci U S A 83:8122–6.

38. Kanai Y, Komoto S, Kawagishi T, Nouda R, Nagasawa N, Onishi M, Matsuura Y, Taniguchi K, Kobayashi T. 2017. Entirely plasmid-based reverse genetics system for rotaviruses. Proc Natl Acad Sci U S A 114:2349–2354.

39. Komoto S, Kanai Y, Fukuda S, Kugita M, Kawagishi T, Ito N, Sugiyama M, Matsuura Y, Kobayashi T, Taniguchi K. 2017. Reverse genetics system demonstrates that rotavirus nonstructural protein NSP6 is not essential for viral replication in cell culture. J Virol 91:e00695–17.

40. Papa G, Venditti L, Arnoldi F, Schraner EM, Potgieter C, Borodavka A, Eichwald C, Burrone OR. 2019. Recombinant rotaviruses rescued by reverse genetics reveal the role of NSP5 hyperphosphorylation in the assembly of viral factories. J Virol 94:e01110–19.

41. Papa G, Venditti L, Braga L, Schneider E, Giacca M, Petris G, Burrone OR. 2020. CRISPR-Csy4-mediated editing of rotavirus double-stranded RNA genome. Cell Rep 32:108205.

42. Jiménez-Zaragoza M, Yubero MP, Martín-Forero E, Castón JR, Reguera D, Luque D, de Pablo PJ, Rodríguez JM. 2018. Biophysical properties of single rotavirus particles account for the functions of protein shells in a multilayered virus. Elife 7:e37295.

43. Dormitzer PR, Greenberg HB, Harrison SC. 2000. Purified recombinant rotavirus VP7 forms soluble, calcium-dependent trimers. Virology 277:420–8.

44. Shaw RD, Vo PT, Offit PA, Coulson BS, Greenberg HB. 1986. Antigenic mapping of the surface proteins of rhesus rotavirus. Virology 155:434–51.

45. Eichwald C, Arnoldi F, Laimbacher AS, Schraner EM, Fraefel C, Wild P, Burrone OR, Ackermann M. 2012. Rotavirus viroplasm fusion and perinuclear localization are dynamic processes requiring stabilized microtubules. PLoS One 7:e47947.

46. Eichwald C, Rodriguez JF, Burrone OR. 2004. Characterization of rotavirus NSP2/NSP5 interactions and the dynamics of viroplasm formation. J Gen Virol 85:625–34.

47. Eichwald C, De Lorenzo G, Schraner EM, Papa G, Bollati M, Swuec P, de Rosa M, Milani M, Mastrangelo E, Ackermann M, Burrone OR, Arnoldi F. 2018. Identification of a small molecule that compromises the structural integrity of viroplasms and rotavirus double-layered particles. J Virol 92:e01943–17.

48. Kroschwald S, Alberti S. 2017. Gel or die: phase separation as a survival strategy. Cell 168:947–948.

49. Lin Y, Mori E, Kato M, Xiang S, Wu L, Kwon I, McKnight SL. 2016. Toxic PR poly-dipeptides encoded by the C9orf72 repeat expansion target LC domain polymers. Cell 167:789–802.e12.

50. Geiger F, Papa G, Arter WE, Acker J, Saar KL, Erkamp N, Qi R, Bravo J, Strauss S, Krainer G, Burrone OR, Jungmann R, Knowles TPJ, Engelke H, Borodavka A. Rotavirus replication factories are complex ribonucleoprotein condensates. bioRxiv https://doi.org/10.1101/2020.12.18.423429

51. Déctor MA, Romero P, López S, Arias CF. 2002. Rotavirus gene silencing by small interfering RNAs. EMBO Rep 3:1175–80.

52. Trask SD, Dormitzer PR. 2006. Assembly of highly infectious rotavirus particles recoated with recombinant outer capsid proteins. J Virol 80:11293–304.

53. Herrmann T, Torres R, Salgado EN, Berciu C, Stoddard D, Nicastro D, Jenni S, Harrison SC. 2021. Functional refolding of the penetration protein on a non-enveloped virus. Nature 590:666–670.

54. Abdelhakim AH, Salgado EN, Fu X, Pasham M, Nicastro D, Kirchhausen T, Harrison SC. 2014. Structural correlates of rotavirus cell entry. PLoS Pathog 10:e1004355.

55. Salgado EN, Garcia Rodriguez B, Narayanaswamy N, Krishnan Y, Harrison SC. 2018. Visualization of calcium ion loss from rotavirus during cell entry. J Virol 92:e01327–18.

56. Salgado EN, Upadhyayula S, Harrison SC. 2017. Single-particle detection of transcription following rotavirus entry. J Virol 91: e00651–17.

57. Trask SD, Kim IS, Harrison SC, Dormitzer PR. 2010. A rotavirus spike protein conformational intermediate binds lipid bilayers. J Virol 84:1764–70.

58. Vesikari T, Clark HF, Offit PA, Dallas MJ, DiStefano DJ, Goveia MG, Ward RL, Schödel F, Karvonen A, Drummond JE, DiNubile MJ, Heaton PM. 2006. Effects of the potency and composition of the multivalent human-bovine (WC3) reassortant rotavirus vaccine on efficacy, safety and immunogenicity in healthy infants. Vaccine 24:4821–9.

59. Ruiz-Palacios GM, Pérez-Schael I, Velázquez FR, Abate H, Breuer T, Clemens SC, Cheuvart B, Espinoza F, Gillard P, Innis BL, Cervantes Y, Linhares AC, López P, Macías-Parra M, Ortega-Barría E, Richardson V, Rivera-Medina DM, Rivera L, Salinas B, Pavía-Ruz N, Salmerón J, Rüttimann R, Tinoco JC, Rubio P, Nuñez E, Guerrero ML, Yarzábal JP, Damaso S, Tornieporth N, Sáez-Llorens X, Vergara RF, Vesikari T, Bouckenooghe A, Clemens R, De Vos B, O’Ryan M, Group HRVS. 2006. Safety and efficacy of an attenuated vaccine against severe rotavirus gastroenteritis. N Engl J Med 354:11–22.

60. Ito N, Takayama-Ito M, Yamada K, Hosokawa J, Sugiyama M, Minamoto N. 2003. Improved recovery of rabies virus from cloned cDNA using a vaccinia virus-free reverse genetics system. Microbiol Immunol 47:613–7.

61. Guglielmi KM, McDonald SM, Patton JT. 2010. Mechanism of intraparticle synthesis of the rotavirus double-stranded RNA genome. J Biol Chem 285:18123–8.

62. Arnold M, Patton JT, McDonald SM. 2009. Culturing, storage, and quantification of rotaviruses. Curr Protoc Microbiol Chapter 15:Unit 15C.3.

63. Yusa K, Zhou L, Li MA, Bradley A, Craig NL. 2011. A hyperactive piggyBac transposase for mammalian applications. Proc Natl Acad Sci U S A 108:1531–6.

64. Eichwald C, Vascotto F, Fabbretti E, Burrone OR. 2002. Rotavirus NSP5: mapping phosphorylation sites and kinase activation and viroplasm localization domains. J Virol 76:3461–70.

65. Sun M, Giambiagi S, Burrone O. 1997. [VP4 protein of simian rotavirus strain SA11 expressed by a baculovirus recombinant]. Zhongguo Yi Xue Ke Xue Yuan Xue Bao 19:48–53.

66. Arnoldi F, Campagna M, Eichwald C, Desselberger U, Burrone OR. 2007. Interaction of rotavirus polymerase VP1 with nonstructural protein NSP5 is stronger than that with NSP2. J Virol 81:2128–37.

67. Buttafuoco A, Michaelsen K, Tobler K, Ackermann M, Fraefel C, Eichwald C. 2020. Conserved Rotavirus NSP5 and VP2 Domains Interact and Affect Viroplasm. J Virol 94.

68. Schindelin J, Arganda-Carreras I, Frise E, Kaynig V, Longair M, Pietzsch T, Preibisch S, Rueden C, Saalfeld S, Schmid B, Tinevez JY, White DJ, Hartenstein V, Eliceiri K, Tomancak P, Cardona A. 2012. Fiji: an open-source platform for biological-image analysis. Nat Methods 9:676–82.

69. Eichwald C, Ackermann M, Nibert ML. 2018. The dynamics of both filamentous and globular mammalian reovirus viral factories rely on the microtubule network. Virology 518:77–86.

70. Eichwald C, Ackermann M, Fraefel C. 2020. Mammalian orthoreovirus core protein μ2 reorganizes host microtubule-organizing center components. Virology 549:13–24.

71. Glück S, Buttafuoco A, Meier AF, Arnoldi F, Vogt B, Schraner EM, Ackermann M, Eichwald C. 2017. Rotavirus replication is correlated with S/G2 interphase arrest of the host cell cycle. PLoS One 12:e0179607.

72. Bauer M, Smith GP. 1988. Filamentous phage morphogenetic signal sequence and orientation of DNA in the virion and gene-V protein complex. Virology 167:166–75.

